# An integration-free CRISPR platform for genetic dissection of respiratory virus host dependencies in human nasal epithelium

**DOI:** 10.64898/2026.10.06.756366

**Authors:** Sophie Wilhelmina van Leur, Julie Tung Sem Chu, Xuesheng Wu, Akanksha Rawat, Nicholas Kim-Wah Yeo, Trena Wan Xuan Lee, Marcus G Mah, Chen Yung-Jiun, Clara Shi Ya Ng, Camille Trupin, Alan Chen-Yu Hsu, Gavin JD Smith, Yaw Shin Ooi, Mart Matthias Lamers

## Abstract

Respiratory viruses, including coronaviruses, generally initiate infection in the upper respiratory tract, where mechanistic studies of host-virus interactions require models amenable to genetic manipulation. Human nasal air-liquid interface (ALI) cultures are realistic models of the nasal epithelium, but robust CRISPR/Cas9 engineering workflows for this primary cell system have been lacking. Here, we established an integration-free CRISPR/Cas9 workflow for genetic engineering of human nasal basal cells, which are subsequently differentiated into gene-edited ALI cultures. GFP-based sorting of RNP-transfected nasal basal cells substantially increases editing, achieving knockout efficiencies of 70–90% across gene targets (B2M, SLC35A1, ACE2, and TMPRSS2). Genetic disruption of ACE2 markedly reduces infection by SARS-CoV-2 and hCoV-NL63, confirming its important role for both viruses in the human nasal epithelium. TMPRSS2 knockout substantially reduces replication of both viruses, demonstrating SARS-CoV-2 depending on TMPRSS2 in the human nasal epithelium and providing genetic evidence in primary human cell cultures that hCoV-NL63 relies on TMPRSS2. This platform enables systematic genetic perturbation in a physiologically relevant model of the primary site of respiratory virus infection, validates ACE2 and TMPRSS2 as targets for medical countermeasures against SARS-CoV-2 and hCoV-NL63, and provides a versatile approach to identify host dependency factors for both circulating and emerging respiratory viruses.

## Main

Respiratory virus infections cause substantial global morbidity and mortality, yet medical countermeasures remain unavailable for most causative agents. In addition, the COVID-19 pandemic, caused by Severe Acute Respiratory Syndrome Coronavirus 2 (SARS-CoV-2), has demonstrated that new respiratory threats can emerge unexpectedly, underscoring the need to improve pandemic preparedness.

Most respiratory viruses are thought to initiate infection in the upper respiratory tract, such as the nasal epithelium, which functions as a critical barrier to infection. This epithelium is therefore an attractive target for prophylactic and therapeutic interventions. Host-targeted strategies to combat virus infections have the potential to offer broad protection against a wide range of viruses, including future pandemic threats, but their development has been limited by gaps in host biology [1]. Defining the host dependency factors that govern viral susceptibility at the primary infection site can thus accelerate the rational development of medical countermeasures. For example, TMPRSS2, a type II transmembrane serine protease exploited by a wide range of respiratory viruses for cell entry, represents an attractive broad-spectrum host target [2–11].

CRISPR-mediated gene editing in cancerous or transformed cell lines has generated a rich catalog of putative host factors, particularly for SARS-CoV-2 [1, 12–16]. However, most of these factors have been identified in cell lines that do not represent the cells targeted in vivo, whereas models that do represent them are not readily amenable to genetic engineering. Accelerating antiviral discovery therefore benefits from CRISPR-editable, physiologically relevant human model systems that faithfully recapitulate the primary site of infection.

In recent years, human nasal airway organoids or air-liquid interface (ALI) cultures have emerged as promising models that closely mimic the human nasal airway epithelium [17–19]. Derived from healthy donor human nasal basal stem cells, they recapitulate key features of the native airway epithelium upon differentiation, including multiciliated cells, secretory lineages, and functional mucus barriers. However, few studies have used CRISPR-mediated gene editing in these cultures due to inefficient gene delivery and the limited capacity of basal cells to proliferate and clonally expand [20–25]. Each of these studies has relied on lentiviral transduction to deliver Cas9 and guide RNAs, which has several drawbacks, including the risk of insertional mutagenesis, the need for prior construct cloning, and, in some cases, prolonged expression of Cas9 and guide RNAs.

In this study, we have developed an integration-free CRISPR/Cas9 workflow in human adult-derived nasal basal stem cells that, after differentiation at ALI, enables functional validation of host dependency factors. We validate this workflow by perturbing host dependency factors for SARS-CoV-2 and human coronavirus NL63 (hCoV-NL63), providing a physiologically relevant platform for identifying therapeutic targets against current and future respiratory viruses.

## Results

### Integration-free CRISPR/Cas9 workflow with eGFP-based sorting enables efficient gene editing in human nasal basal cells

Although traditionally human nasal basal cells could only be passaged 1-3 times before becoming senescent, recent studies using defined culture medium have substantially extended the proliferative capacity in two-dimensional (2D) culture using extracellular matrix (ECM)-coated culture vessels and a defined medium [18, 19, 26]. In our hands, these cultures can be passaged 8-12 times before their proliferative and differentiation capacity declines. We used these proliferative 2D-expanded adult-derived human nasal basal cells to establish a CRISPR-editing platform for dissecting host dependency factors in the nasal epithelium.

We developed an integration-free Ribonucleoprotein (RNP)-based CRISPR/Cas9 knockout workflow in which cells are CRISPR-edited as basal cells, expanded as pools, and then differentiated at ALI (**Fig. 1a-b)**. Knockout pools rather than clonal lines are used because the extensive passaging required for clonal expansion can exceed the passaging limit of nasal basal cells. Briefly, Cas9 nuclease/gRNA RNP complexes were transiently transfected alongside *eGFP* mRNA to enable fluorescence-based sorting of transfected cells. Sorted cell pools were expanded and sequenced to verify knockout efficiency. The workflow was initially validated using an sgRNA targeting the invariant light chain of the HLA class I proteins, beta-2 microglobulin (B2M) [27]. Targeting B2M depletes surface HLA-I expression, allowing knockout efficiency to be assessed by flow cytometry using an anti-HLA-I antibody. As shown in **Figure 1c**, 47.1% of cells transfected with B2M sgRNA RNP yielded lost HLA-I expression, whereas 100% of the non-targeting control sgRNA cells retain HLA-I expression. Enriching for eGFP expression increased the knockout efficiency to 86.8%. In all downstream experiments, we used this sorting-based method to enrich for knockout cells.

**Fig. 1.**
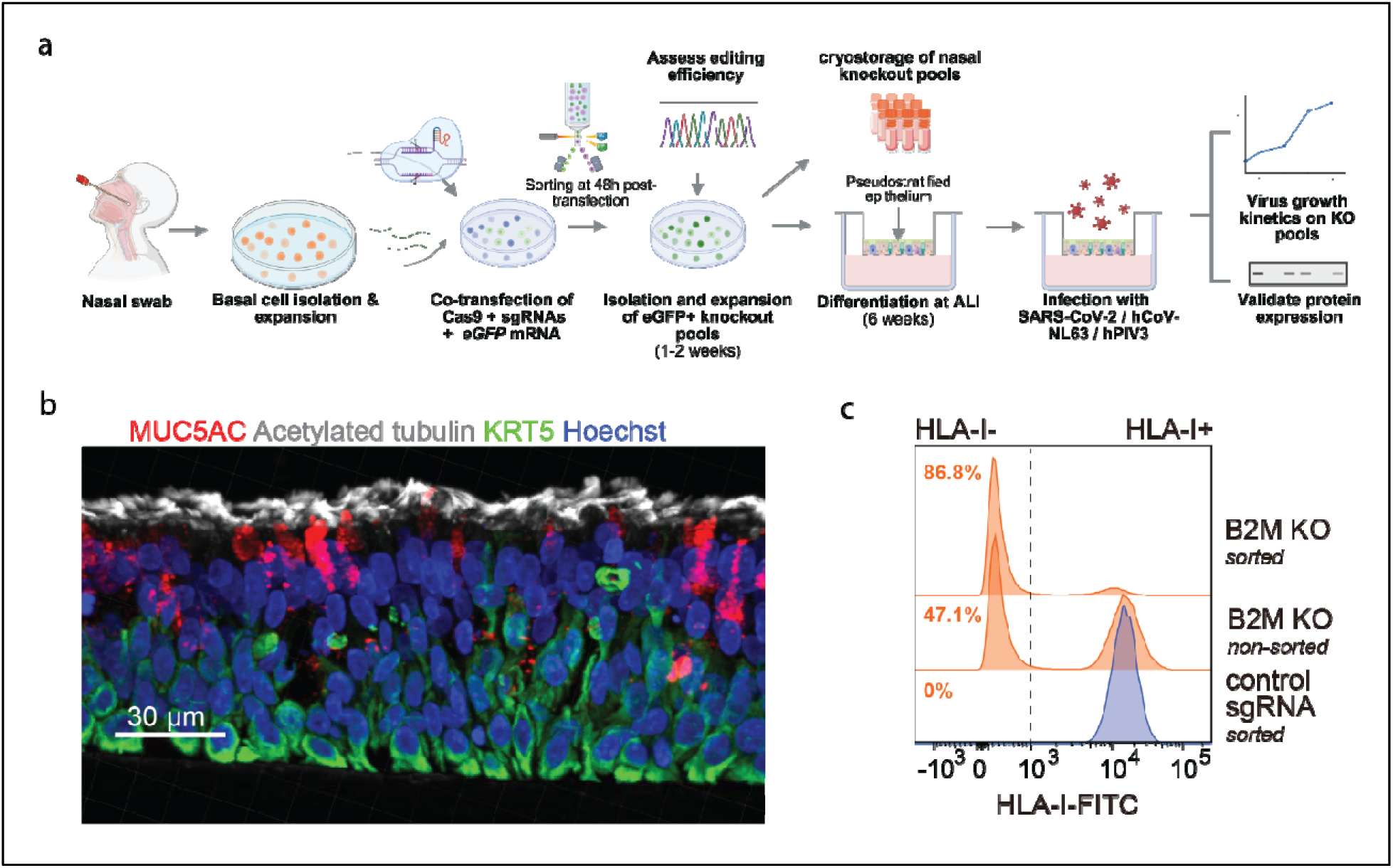
An integration-free CRISPR/Cas9 workflow enables efficient gene editing in nasal basal cells. **a** Schematic of the human adult-derived nasal basal CRISPR/Cas9 workflow. Created using BioRender.com. Illustrations are conceptual and not derived from experimental data. Basal cells from healthy donor nasal brushings were expanded and differentiated at ALI to form a pseudostratified epithelium. For gene editing, basal cells were transfected with Cas9 protein, sgRNA, and *eGFP* mRNA to allow for fluorescence-based sorting of transfected cells at 48 hours post-transfection. Editing efficiency is validated by Sanger sequencing and/or flow cytometry prior to cryostorage and ALI differentiation or by western blotting after differentiation. Mature knockout and non-targeting control sgRNA cultures are infected with SARS-CoV-2, hCoV-NL63 or hPIV3. Viral replication is assessed by fluorescence, confocal imaging and RT-qPCR. **b** Representative cryosection of a nasal ALI culture after 6 weeks of differentiation, stained for ciliated cells (acetylated tubulin), goblet cells (MUC5AC), basal cells (KRT5), and nuclei (Hoechst), acquired at 40× on the Andor BC43. **c** Representative flow cytometric analysis of B2M knockout with and without sorting (GFP)-based enrichment for transfected cells, as measured by loss of HLA-I expression at 14 days post-transfection. Data in **b** and **c** are representative of at least 3 independent experiments.

### SLC35A1 knockout in nasal basal cells validates CRISPR editing workflow with reduced hPIV3 susceptibility

As proof-of-concept, the CRISPR/Cas9 workflow was further validated by knocking out solute carrier family 35 member A1 (SLC35A1), which encodes the only known transporter for sialic acids into the Golgi [28]. SLC35A1 was selected as it offers an additional facile means of assessing editing efficiency by flow cytometry, using lectin staining to detect loss of terminal sialylation, and further allows validation of the workflow in an infection-based assay using a reporter virus known to depend on terminal sialic acids for entry, Human respirovirus 3, commonly known as Human parainfluenza virus 3 (hPIV3) [29]. Knocking out SLC35A1 led to a near complete loss of surface sialylation (**Supplementary Fig. 1a**). We obtained a knockout efficiency of 83%, as assessed by flow cytometric analysis of *Sambucus Nigra Agglutinin* (SNA) lectin staining, confirming loss of terminal sialylation in SLC35A1 knockout cells. These cells were differentiated for 3 weeks at ALI prior to infection with hPIV3-eGFP at 100,000 FFU/well. Live imaging of eGFP signal at 72 hours post-infection (h.p.i.) showed visibly reduced hPIV3-eGFP infection in SLC35A1 knockout ALI cultures compared to non-targeting sgRNA controls (**Supplementary Fig. 1b**). This reduction was confirmed by quantification of eGFP-positive foci at 72 hours post-infection (**Supplementary Fig. 1c**). In addition to the reduced number of infected foci, single-field images of infected regions revealed lower eGFP signal intensity within infected regions of SLC35A1 knockout cultures, suggesting reduced viral replication or spread (**Supplementary Fig. 1d**).

### ACE2 knockout demonstrates the utility of CRISPR editing workflow, abrogating SARS-CoV-2 and reducing hCoV-NL63 infection

Having established the workflow, we next targeted ACE2, the key entry receptor for SARS-CoV-2 and hCoV-NL63 [4, 30, 31]. We included hCoV-NL63, a circulating common human coronavirus that infects the upper respiratory tract and also uses ACE2 for cellular entry, to provide a complementary ACE2-dependent respiratory virus for functional validation of our platform [30, 32]. We knocked ACE2 out in human nasal basal cells, but since ACE2 is not expressed at the basal cell stage, we performed western blotting following 6-week differentiation of the ALI cultures to confirm the knockout. We demonstrated an 83% knockout of ACE2 at the protein level of a representative western blot (**Fig. 2a-b)**. In two independent batches of ACE2 knockout ALI cultures generated from the same nasal donor, we observed abrogated SARS-CoV-2-mCherry infection by RT-qPCR (**Fig. 2c, Supplementary Fig. 2a**). Live-imaging further demonstrated that, at 72 hours post-infection, an average of 25% of the surface area was infected in non-targeting control cultures, compared to only 0.02% in ACE2 knockout cultures (**Fig. 2e-f**). In addition, infection of the same set of ACE2 knockout cultures with hCoV-NL63 demonstrated that ACE2 knockout also markedly reduces hCoV-NL63 infection, as seen by a >90% reduction in copy numbers at 48 hours post-infection (**Fig. 2d and Supplementary Fig. 2b**). These results confirm the critical role for ACE2 in SARS-CoV-2 and hCoV-NL-63 infection in the human nasal epithelium.

**Fig. 2.**
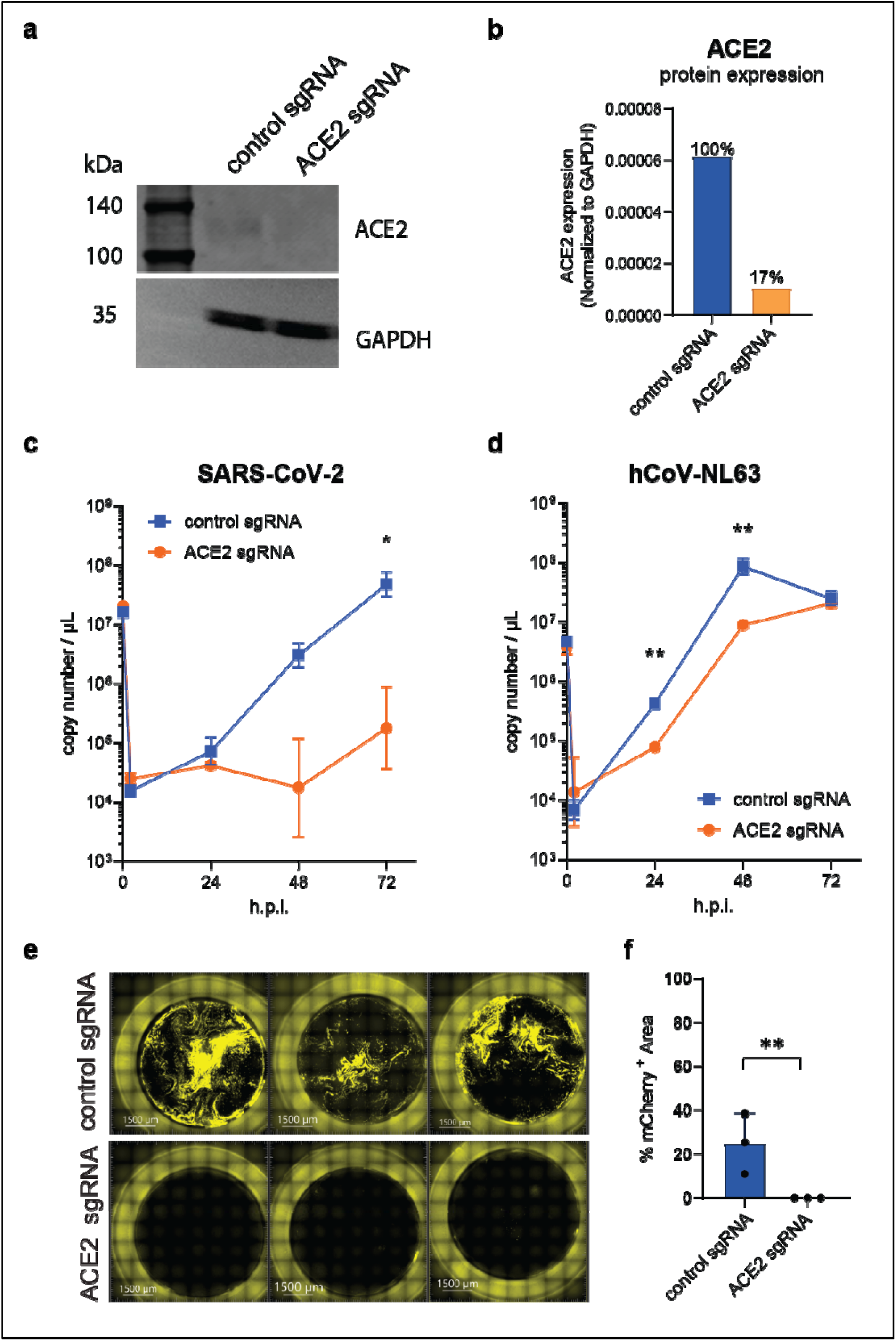
ACE2 knockout blocks SARS-CoV-2-mCherry and significantly reduces hCoV-NL63 infection in nasal ALI cultures. **a** A representative western blot confirms ACE2 protein reduction in ACE2 knockout (ACE2 sgRNA) vs. non-targeting sgRNA (control sgRNA). **b** Western blot quantification indicates ∼83% knockout after sorting and differentiation. **c-d** Infection of nasal ALI cultures with **c** SARS-CoV-2-mCherry and **d** hCoV-NL63 (both 1,000 FFU/well) demonstrate markedly reduced viral RNA in the ACE2 knockout (RT-qPCR, triplicate ALI cultures). Statistical significance was assessed using two-way ANOVA followed by Holm-Šidák’s multiple-comparisons test, comparing the knockout with the control at each time point with * p < 0.05 and ** p<0.01. **e** Live imaging of SARS-CoV-2-mCherry infection at 72 hours post-infection (h.p.i.) shows minimal infection in ACE2 knockout (561 nm excitation, yellow epifluorescence channel, 10x, 7×7 stitched grid). **f** Area-based quantification of mCherry fluorescence at 72 h.p.i. confirms near complete loss of infection. The percentage of mCherry-positive area per well was quantified in ImageJ from three ALI cultures per condition. Statistical differences between sgRNA control and target knockout were tested using Welch’s unpaired t-test on log₁₀-transformed data. Data are shown as mean ± SD (raw %).

### TMPRSS2 is required for SARS-CoV-2 and hCoV-NL63 infection in nasal ALI cultures

We next investigated the role of TMPRSS2 in SARS-CoV-2 and hCoV-NL63 entry in the nasal epithelium. It was previously established that SARS-CoV-2 relies on serine protease-dependent entry in airway ALI cultures, whereas in some cell lines endosomal cathepsins are used [4, 33–36]. These studies rely on treatments with non-specific protease inhibitors, like camostat for serine proteases including TMPRSS2, and E64D for cathepsins.

First, we verified that both SARS-CoV-2 and hCoV-NL63 entry in the nasal ALI model was dependent on serine protease-mediated entry rather than late endosomal entry. Briefly, unedited human nasal ALI cultures were pre-treated with E64D or camostat for 1 hour, to block each entry pathway (**Fig. 3a**). Inhibitors were maintained throughout subsequent washing steps and apical harvests, to ensure continuous inhibition of the targeted entry pathway for the duration of the experiment. We demonstrate that endosomal cathepsins do not contribute to viral infection in this model, as E64D treatment had no effect on virus replication. In contrast, both SARS-CoV-2 and hCoV-NL63 relied heavily on serine proteases for infection, with camostat treatment resulting in a complete loss of SARS-CoV-2 replication and a substantial reduction in hCoV-NL63 replication (**Fig. 3b-d, Supplementary Fig. 3a-b**). Next, we investigated whether this serine protease dependence is specifically attributable to TMPRSS2 by generating TMPRSS2 knockout nasal cells. The knockout was generated using a pool of three TMPRSS2-targeting guides, as single guides yielded variable knockout efficiency for this gene across batches (data not shown). Sanger sequencing confirmed that our sorted knockout pools achieved approximately 80% editing efficiency, as shown by representative ICE traces (**Supplementary Fig. 4a-c**). This translated into a functional knockout, with representative western blot analysis of differentiated ALI cultures showing a 73% reduction in the active, cleaved form of TMPRSS2 (**Fig. 3e-f**). Next, we infected TMPRSS2 knockout and control nasal ALI cultures with SARS-CoV-2-mCherry (**Fig. 3g**). TMPRSS2 knockout resulted in a marked reduction in SARS-CoV-2-mCherry infection, with imaging revealing a near-complete loss of infection, limited to only a few isolated infection foci (**Fig. 3i-k**).

**Fig. 3.**
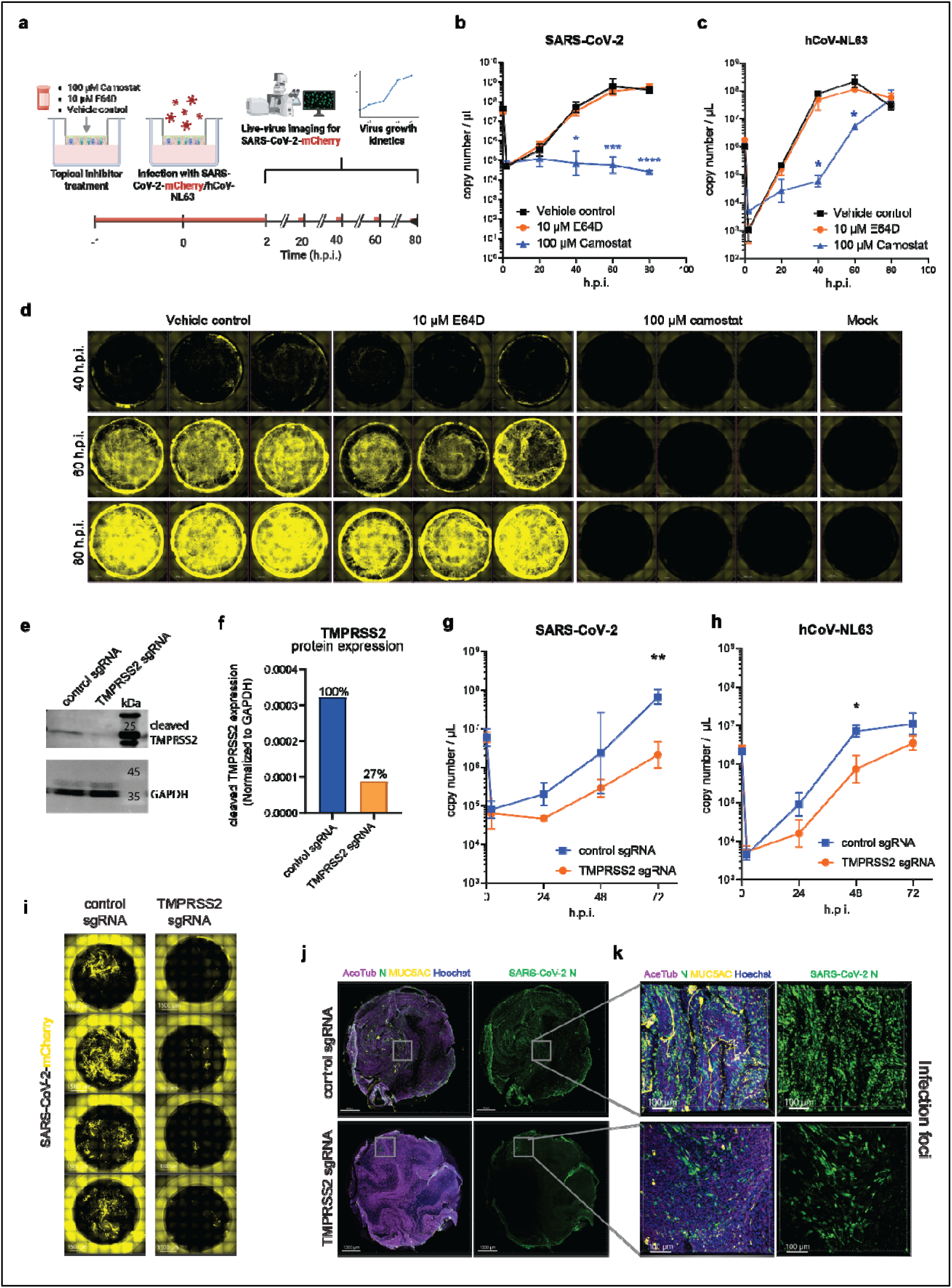
TMPRSS2 knockout reduces SARS-CoV-2-mCherry and hCoV-NL63 infection in nasal ALI cultures. **a** Schematic of the inhibitor treatment workflow. Created using BioRender.com. Illustrations are conceptual and not derived from experimental data. **b** SARS-CoV-2-mCherry and **c** hCoV-NL63 Infection of nasal ALI cultures (1,000 FFU/well) show significantly reduced viral RNA following pre-treatment with 100 µM camostat, but not E64D (E gene RT-qPCR; n = 1 donor, triplicate ALI cultures). **d** Live-imaging at 40, 60, and 80 h.p.i. shows minimal SARS-CoV-2-mCherry infection in camostat-treated nasal ALI cultures (561 nm excitation, yellow epifluorescence channel, 10x, 7x7 stitched grid). **e** Representative western blot confirms reduction in cleaved TMPRSS2 levels in the TMPRSS2 knockout (TMPRSS2 sgRNA) vs. non-specific control sgRNA (control sgRNA). **f** Western blot quantification of the active, cleaved form of TMPRSS2 shows ∼73% knockout after sorting and differentiation. **g** Infection of nasal ALI knockout cultures with **g** SARS-CoV-2-mcherry or **h** hCoV-NL63 (1,000 FFU/well) demonstrate significantly reduced viral RNA in TMPRSS2 knockout (E gene RT-qPCR, **g** quadruplicate and **h** triplicate ALI cultures). **i** Live imaging at 72 hours post-infection (h.p.i.) with SARS-CoV-2-mCherry shows minimal infection in TMPRSS2 knockout (561 nm excitation, yellow epifluorescence channel, 10x, 7x7 stitched grid). **j** Differentiated ALI cultures retain ciliated (Acetylated Tubulin (AceTub)) and goblet (MUC5AC) cells, but TMPRSS2 knockout cultures exhibit markedly reduced infection by SARS-CoV-2 Nucleocapsid protein (N) staining (Confocal, 10x), with **k** residual infection restricted to isolated foci (20× objective). Statistical significance was assessed using two-way ANOVA followed by **b,c,h** Dunnett’s or **g** Holm-Šidák’s multiple-comparisons test, comparing each treatment or knockout with the control at each time point with * p < 0.05 and ** p<0.01, *** p<0.001 and **** p<0.0001 (for **h** hCoV-NL63, the TMPRSS2 and ACE2 sgRNA (**Supplementary Fig. 2b**) conditions were generated in the same knockout batch and therefore share the same non-targeting control data).

To confirm that this phenotype is consistent and not donor-dependent, we generated TMPRSS2 knockout ALI cultures once more in the same donor (**Supplementary Fig. 5**) and in an independent donor (**Supplementary Fig. 6a-d).** Again, SARS-CoV-2-mCherry infection was observed to be significantly reduced in this same donor, whereas productive replication was completely abrogated in the alternative donor. Together, these results demonstrate that the requirement for TMPRSS2 in SARS-CoV-2 entry is consistent across donors.

Finally, we assessed whether hCoV-NL63 infection similarly relies particularly on TMPRSS2 for viral entry. Indeed, TMPRSS2 knockout reduced hCoV-NL63 infection to a comparable extent as the knockout of the entry receptor ACE2, indicating that TMPRSS2 is also a critical host factor for hCoV-NL63 (**Fig 3h, Supplementary Fig 2b, Figure 2d)**.

## Discussion

In biology, a model is of limited use if it cannot be genetically manipulated efficiently and reliably. Here, we developed an integration-free CRISPR/Cas9-edited nasal ALI platform that enables functional interrogation of host dependency factors for respiratory viruses. We demonstrated that this workflow efficiently and reliably generated functional knockout pools which could be differentiated at ALI and utilized to dissect host dependency factors.

A key feature of this platform is that efficient genome editing can be achieved in primary nasal basal cells without the use of integrating viral vectors or clonal expansion. Using B2M knockout as an initial proof-of-concept target, we showed that eGFP-based sorting of transfected cells substantially improved editing efficiency in nasal cells, increasing knockout frequency from 47.1% to 86.8%, as measured by loss of surface HLA-I expression. This is comparable to editing efficiencies reported by Rapiteanu and colleagues [27], who achieved highly efficient B2M knockout (90–97%) in bronchial basal cells using RNP lipofection or electroporation. Whereas that study achieved high transfection efficiency without sorting, efficient editing of the nasal cells used here required sorting to raise editing rates and obtain consistent results. Across the genes targeted in this study, editing efficiencies were consistently high (approximately 70–90%) at the DNA and protein level. Notably, the maintenance of high editing efficiency after differentiation indicates that this workflow yields phenotypically informative ALI cultures suitable for downstream functional assays. ACE2 knockout ALI cultures were resistant to SARS-CoV-2 and showed significantly reduced hCoV-NL63 infection, confirming ACE2 as the primary receptor for both viruses in the human nasal epithelium [4, 31, 32].

Previous studies also demonstrated that SARS-CoV-2 entry relies on serine proteases in nasal ALI cultures by showing that camostat effectively blocked infection [33, 34]. However, broad-spectrum inhibitors such as camostat cannot resolve individual protease usage. Overexpression assays in cell lines, in turn, overestimate the range of proteases a protein can use, further complicating protease usage dissection studies [2, 4]. We previously tackled this using CRISPR-edited clonal intestinal organoids and infecting them with infectious virus [2], but clonal outgrowth infeasible in airway cultures. Here, we have overcome this caveat and enabled specific host factor dissection in nasal ALI knockout pool cultures using RNP-based CRISPR gene editing and sorting to increase editing efficiency. TMPRSS2 deficiency strongly decreased infection, confirming that TMPRSS2 is indeed the main priming protease for SARS-CoV-2 in the human nasal epithelium. This specificity in protease usage is consistent with our previous study using CRISPR-edited intestinal organoids, which identified TMPRSS2, but not other type II transmembrane proteases, as a critical factor for SARS-CoV-2 infection in the gut [2].

We also provide evidence that hCoV-NL63 relies predominantly on serine protease-dependent entry, not endosomal entry, in the nasal epithelium, supporting the results found in bronchial ALI cultures [32, 38]. As with SARS-CoV-2 and hCoV-229E [39, 40], cell lines appear to misidentify endosomal entry as the pathway used by hCoV-NL63, indicating that physiologically relevant models are required to accurately identify host dependency factors. We further show that this serine protease dependence is attributable specifically to TMPRSS2, as knockout of this gene significantly reduced hCoV-NL63 infection. To our knowledge, this is the first genetic evidence, in any model system, that hCoV-NL63 depends on TMPRSS2. It remains biologically surprising that coronaviruses primarily appear to rely specifically on TMPRSS2, given the wide range of serine proteases expressed in the human airway and intestinal epithelium [2, 41].

Future work should extend this investigation to other SARS-CoV-2 strains, particularly Omicron variants. Previous studies have debated whether these variants have adapted to rely less on cell surface entry, or specifically TMPRSS2, and whether this shift explains their preferential infectivity in the upper rather than lower respiratory tract [33, 34, 42, 43]. It should be investigated whether Omicron has adapted to specifically utilize other proteases instead of TMPRSS2 in the nasal and alveolar epithelium to underpin the mechanistic basis of this altered entry phenotype and potentially refine the development of medical countermeasures. In addition, this platform can be readily used for host factor identification when new pandemic threats emerge.

While our integration-free workflow offers several advantages over lentiviral, clonal approaches, it also has notable limitations. First, the current approach is not readily scalable for high-throughput applications. Generating validated knockout ALI cultures is both costly and time-intensive, requiring approximately eight weeks from transfection to infection readout. Secondly, as CRISPR/Cas9 delivery is performed at the basal stem cell stage, our approach is limited to genes not involved in basal cell proliferation or differentiation. Other limitations of our workflow include the inability to generate complete knockouts and donor-to-donor variation in editing efficiency, with the latter exemplified by the TMPRSS2 knockout, which yielded abrogated infection in one donor but reduced infection in another. This variability might also be an advantage, as these differences likely reflect real-world biology (e.g., donor-to-donor variation in the expression of compensatory serine proteases) affecting the success of therapeutic interventions. A second limitation of the pooled approach is that residual infection of unedited cells makes it difficult to distinguish essential host factors from those that are merely required for efficient replication. In intestinal organoids, complete knockouts can be generated using established CRISPR/Cas9 workflows, as intestinal organoids can be expanded clonally. Conversely, despite the recent advancements in medium composition, the extensive passaging required for clonal expansion, is still incompatible with the current human nasal basal cell models [19]. Nevertheless, the workflow identified host dependency factors for all viruses tested, indicating that it can be applied to other respiratory viruses that infect these cultures.

In conclusion, our findings establish the feasibility of dissecting host dependency factors in nasal ALI cultures and underscore the critical roles of ACE2 and TMPRSS2 in SARS-CoV-2 and hCoV-NL63 infection. By enabling systematic genetic perturbation in a physiologically relevant model of the primary site of respiratory virus infection, this approach provides a versatile platform for identifying and validating host dependency factors of circulating and emerging respiratory viruses.

## Methods

### Human nasal basal cell culture

Human adult nasal cells were obtained from healthy adult donors via brushing of the nasal cavity using an interdental brush (Curaprox, CPS 06 PRIME START,) after informed consent. All donor materials were anonymized. The study protocol was approved by the NUS Institutional Review Board (NUS-IRB-2021-706). Isolation and expansion of human nasal stem cells followed a protocol adapted from Breugem et al. (2024) and Amatngalim et al. (2022) [19, 44]. In brief, the nasal brushings were immediately placed into collection medium (Advanced DMEM/F12 with 1× GlutaMax and 1× HEPES, supplemented with 100 µg/mL Primocin (InvivoGen, ant-pm-2) and 5 µM Y-27632 (MedChemExpress, HY-10583)) and stored on ice. Following centrifugation, collection medium was removed, and brushes were incubated in TrypLE (Gibco, 12604-013) containing 5 µM Y-27632 for 5–10 min at 37°C to dissociate cells, with vigorous agitation until brushes appeared visibly clean. Cells were quenched and washed from brushes using collection medium, pooling all supernatant until no cells remained visible on the brush by light microscopy. Cells were pelleted (400 × *g*, 3 min at room temperature) and washed with collection medium, with additional washes performed as needed to remove excess mucus or debris. Cells were either cryopreserved or directly seeded into expansion medium (**Table 1**) with iMatrix-511 SILK (1µg/cm^2^; REPROCELL, NP892-021). Cells were passaged 1:10 every 7-10 days upon reaching 80-90% confluency.

**Table 1.** Human nasal expansion medium as first described by Amatngalim et al. [. **26].**

| Reagent | Concentration | Supplier | Cat No. |
| --- | --- | --- | --- |
| **BEpiCM-b with 50 µg/mL Primocin, 1× GlutaMax** | 50% (v/v) | ScienCell | 3211 |
| **Advanced DMEM/F12 with 50 µg/mL Primocin, 1× GlutaMax and 1× HEPES** | 44.8% (v/v) | Gibco | 12634-034 |
| **B27 supplement** | 2% (v/v) | Gibco | 17504-44 |
| **Nicotinamide** | 5 mM | Sigma | N0636 |
| **N-Acetylcysteine** | 1.25 mM | Sigma | A9165 |
| **DAPT (NOTCH inhibitor)** | 5 µg/mL | MedChemExpress | HY-13027(10 mg) |
| **FGF 10** | 100 ng/mL | Peprtech | 100-26-100 |
| **R-spondin1** | 100 ng/mL | IpA Therapeutics | R006 |
| **Y-27632 (ROCK inhibitor)** | 5 µM | MedChemExpress | HY-10583G |
| **FGF 7 / KGF** | 25 ng/mL | Peprtech | 100-19-50 |
| **HGF** | 25 ng/mL | Peprtech | 100-39H-25 |
| **A83-01 (TGF-β inhibitor)** | 1 µM | MedChemExpress | HY-10432 |
| **(±)-Epinephrine hydrochloride** | 0.5 µg/mL | Sigma | E4642-5G |
| **3,3',5-Triiodo-L-thyronine sodium salt** | 100 nM | Sigma | T6397-100MG |
| **DMH-1 (BMP inhibitor)** | 1 µM | MedChemExpress | HY-12273(10 mg) |
| **Hydrocortisone** | 0.5 µg/mL | MedChemExpress | H0888-1G |
| **EGF** | 5 ng/mL | Peprtech | AF-100-15-500 |
| **SB202190 (p38 inhibitor)** | 500 nM | MedChemExpress | HY-10295 |

### Differentiation at ALI

Nasal basal cells were made single cell using TrypLE, followed by washing in AdDF+++ (Advanced DMEM/F12 with 1× Primocin, 1× GlutaMax and 1× HEPES). Transwell inserts were coated with 1:400 Rat Collagen type I (Corning, 354249) in DPBS^-/-^ (without Ca²^+^/Mg²^+^; 1^st^ Base, BUF-2046-1X500ml) for 20 min at 37°C, before seeding nasal basal cells at a density of 150,000 cells per well on a 6.5 mm Transwell (STEMCELL Technologies, 38024). The cells were cultured in a 1:1 mixture of expansion medium and Pneumacult-ALI Medium (STEMCELL Technologies, 05001) until confluent. After 2-4 days, the confluent monolayers were transitioned to ALI in Pneumacult-ALI Medium supplemented with 50 µg/mL Primocin, with medium changes every 3-4 days and apical washes with DPBS^+/+^ (Ca²⁺/Mg²⁺; Gibco, 14040-133) every 3-4 days after 3 weeks of differentiation.

### Virus stock production

For this study, three viruses were utilized; hCoV-NL63/2167M/Tanzania/2018 (PQ7627, passage 3) as well as recombinant mCherry-expressing SARS-CoV-2 (SARS-CoV-2-mCherry passage 4, [45]) and eGFP-expressing Human respirovirus 3 (hPIV3-eGFP, passage 3, [46]). The recombinant SARS-CoV-2-mCherry (generated from Wuhan-1 isolate (NC_045512)) was a kind gift from Dr. Suresh Mahalingam (Griffith University). The virus was rescued as described before [45], with the exception that supernatant from the DNA transfection was directly inoculated onto nasal ALI cultures instead of cell line-based propagation. The recombinant hPIV3-eGFP was a kind gift from Dr. Benhur Lee (Icahn School of Medicine, Mount Sinai). hPIV3-eGFP was rescued using a previously described reverse genetics system [46].

All SARS-CoV-2 strains and hCoV-NL63 were propagated in 6-7-weeks differentiated human nasal ALI cultures. Briefly, ALI cultures on 12 mm Transwell inserts (STEMCELL Technologies, 38023) were refreshed basolaterally with Pneumacult-ALI Medium and washed apically with 200 µL DPBS^+/+^ for 10 min at 37°C prior to infection to remove residual surface mucus. Subsequently, virus inoculum was overlayed onto the ALI at 1,000 FFU/well in 200 µL and incubated at 37°C for 2 hours. This corresponds to an approximate MOI of 0.003, based on an estimated 3.4 × 10⁵ luminally accessible cells, and an MOI of 0.001 based on 1.0 × 10^6^ total cells per 12-well Transwell ALI culture. Afterwards, the nasal ALI cultures were washed twice with AdDF+++, with the second wash being incubated at 37°C for 10 min. The nasal cultures were returned to 37°C for 4 days, with apical harvest on day 1-4. For apical harvest, 300 µL AdDF+++ was added to the apical side and incubated at 37°C for 10 min. Supernatant was gently triturated on the cells, transferred to a 50 mL tube, then stored at 4°C until all harvests were completed. Following day 4 harvest, pooled SARS-CoV-2-mCherry supernatant was centrifuged at 1,000 × *g* for 5 min to remove cell debris, transferred to a new tube and aliquoted. In contrast, pooled hCoV-NL63 harvests were aliquoted directly without centrifugation, as infectious hCoV-NL63 was found to be associated with the cellular fraction (data not shown). The aliquoted virus stocks were stored at -80°C.

Titres were determined for SARS-CoV-2 and hCoV-NL63 by foci forming units (FFU) assay. These stock titres were determined in Calu-3 cells (ATCC, HTB-5). Calu-3 cells were seeded on a 96-well plate at a density of 40,000 cells/well 24 hours prior to infection in Opti-MEM™ + 1× Glutamax + 10% FBS, 1% Pen/Strep (Thermo Fisher, 15140-122,) and 5 µM Y-27632. The culture supernatant was defrosted at room temperature and 10-fold serially diluted in Opti-MEM™ + 1× Glutamax (Gibco, 3259304). Triplicates of each dilution were added to respective wells and incubated for 8 hours before fixation with 10% Neutral-buffered formalin (10% NBF, Sigma Aldrich, HT501128-4L) for 60 min. Afterwards, buffer was exchanged for DPBS^-/-^ and stored at 4°C until staining.

Immunofluorescent staining of titration plates was performed by simultaneous blocking and permeabilizing with blocking buffer containing 3% BSA (Sigma-Aldrich, A9418), 0.1% Triton X-100 (Sigma-Aldrich, T8787) in DPBS^-/-^ for 1 hour at room temperature, followed by incubation with primary antibody diluted in blocking buffer for 2 hours at room temperature. SARS-CoV-2-infected samples were stained with rabbit anti-SARS-CoV-2 nucleocapsid protein antibody (1:5000; SinoBiological, 40143-T62) and hCoV-NL63-infected samples were stained with rabbit anti-hCoV-NL63 nucleocapsid antibody (1:5000; SinoBiological, 40641-T62). Samples were subsequently washed three times for 5 min in blocking buffer, followed by incubation with secondary antibody, donkey anti-rabbit IgG Alexa Fluor™ 488 (1:1000; Thermo Fisher, A-21206) diluted in blocking buffer for 2 hours at room temperature. Hoechst (1 µg/mL; Thermo Fisher, 33342) and Phalloidin-Alexa Fluor Plus™-647 (1:2000, Thermo Fisher, A30107) were added during the final 30 min of secondary antibody incubation. Following three additional 0.1% Triton X-100 in DPBS^-/-^ washes, samples were resuspended in DPBS^-/-^ and stored at 4°C until image acquisition. Image acquisition was done on Andor BC43 benchtop spinning disk confocal microscope (Oxford Instruments). Full-well images were acquired using brightfield, blue-channel (405 nm excitation; 430–480 nm emission), green-channel (488 nm excitation; 500–550 nm emission) and red-channel fluorescence (638 nm excitation; 671-745 nm emission). Images were analyzed using Imaris Spot analysis (version 11, Oxford Instruments) for spot/focus quantification and calculation of FFU/mL titres.

For propagation of hPIV3-eGFP, an epithelial cell line derived from the kidney of adult rhesus monkeys, LLC-MK2 cells were used, that overexpress TMPRSS2. The TMPRSS2-overexpressing LLC-MK2 cells were maintained in DMEM (Gibco, 11965-092) supplemented with 5% FBS (Gibco; A5256801) and infected with hPIV3-eGFP at 50% confluency. The cultures were incubated at 37°C with 5% CO₂ for 2 days. Supernatant was harvested and cleared of debris by spinning at 500 x *g* for 10 min at 4°C before transferring to a new vial and aliquoting for storage at -80°C.

HPIV3-eGFP stock titres were determined by a 50% tissue culture infectious dose (TCID₅₀) assay with TMPRSS2-overexpressing LLC-MK2 cells. Cells were seeded at 5,000 cells per well in 96-well plates containing 100 μL of DMEM supplemented with 2% FBS. Cells were inoculated with 100 μL of 10-fold serial dilutions of virus stock and incubated at 37°C with 5% CO₂. At 7 days post-infection, hPIV3-eGFP infection was identified by monitoring eGFP expression under a fluorescence microscope. TCID₅₀ values were calculated from four replicate wells per dilution using the Spearman–Kärber method.

### CRISPR/Cas9 integration free editing

sgRNAs were designed for human ACE2, TMPRSS2, B2M and SLC35A1 using the CRISPR design tool of Synthego and the OmniGuide RNA Design Tool of Genscript. We included an sgRNA for B2M that was previously published by Sanjana and colleagues (2014) [47]. sgRNAs were ordered as EasyEdit sgRNA through the Genscript website. TrueGuide™ sgRNA Negative Control was used as non-targeting sgRNA control (Invitrogen, A35526). The sgRNAs, their genomic identifiers, and sequencing primers used to validate the knock-out can be found in **Supplementary Table 1 and 2**. To generate knockouts in the genes-of-interest, Cas9-sgRNA RNP complexes were formed using the CRISPRMAX transfection system with Cas9 Plus reagent (Invitrogen, CMAX00008). Briefly, 24 hours prior to transfection, 70,000 cells/well were seeded in a 24-well plate in expansion medium supplemented with iMatrix Silk-511 (1 µg/cm^2^). For transfection, 7.5 pmol of sgRNA (2.5pmol per sgRNA when using the triple sgRNA approach) was mixed with 7.5 pmol TrueCut™ Cas9 Protein v2 (Thermo Fisher, A36498) and 50 ng commercial *eGFP* mRNA (GenScript, RP-A00008) in Opti-MEM™+1× Glutamax, followed by addition of 2.5 µL Cas9 plus reagent and thorough mixing. In a separate tube 1.5 µL CRISPRmax reagent was mixed into Opti-MEM™+1× Glutamax. After 5 min of incubation at room temperature, these reactions were added together and incubated for 10-15 min at room temperature, before drop-wise addition to the cells. Prior to addition of the complexes, the basal cells were refreshed with fresh expansion medium. Afterwards the cells were incubated for 48 hours prior to sorting.

Fluorescence-based sorting was performed at 48 hours post-transfection, coinciding with peak *eGFP* mRNA expression, which is transiently detectable from approximately 24 to 96 hours post-transfection. Basal cells were sorted using an Aria II FACS sorter with the 488 nm laser line. Briefly, the cells were collected by washing once in DPBS^-/-^, dissociating with TrypLE for 5–10 min, and collecting in collection medium. Cells were then washed once by centrifugation with collection medium at 400 × *g* for 5 min, before resuspending in FACS buffer (DPBS^-/-^, 2% FBS, 2 mM EDTA, 5 µM Y-27632). Cells were gated on FSC-A vs. SSC-A to exclude debris, followed by singlet discrimination (FSC-H vs. FSC-A). Within the single-cell gate, eGFP expression was assessed by histogram analysis, setting the positive population using an eGFP only transfected positive control and untransfected negative control. The GFP-positive population was sorted and expanded in expansion medium supplemented with iMatrix Silk-511 (1 µg/cm^2^). Upon reaching confluency (7-10 days), the cells were collected by washing once in DPBS^-/-^, dissociating with TrypLE for 5-10 min and collecting in collection medium. After washing twice with collection medium, the cells were seeded on Transwells for differentiation as described above and collected for genomic DNA isolation for KO validation using Sanger sequencing.

After 3-6 weeks of differentiation, cultures were assessed for knockout efficiency by western blot and were subsequently used for infection experiments (3 weeks hPIV3, 6 weeks coronavirus infections). Confirmation of knockout by western blot was performed after differentiation, as several target proteins, including ACE2, are minimally expressed at the undifferentiated basal cell stage and are upregulated upon differentiation [48].

### Flow cytometry

B2M knockout efficiency was assessed 10-14 days post transfection in the nasal basal cells by flow cytometric analysis of surface HLA-I expression. Culture medium was removed from nasal basal cells, and cells were washed with DPBS^-/-^. Cells were dissociated with 200 µL TrypLE per well (37°C, 5-10 min), quenched with 300 µL AdDF+++, and pelleted (400 × *g*, 5 min). Cells were washed once in FACS buffer (DPBS^-/-^, 2% FBS, 2 mM EDTA) and pelleted again (400 × *g*, 5 min). Cells were resuspended in FACS buffer containing FITC-conjugated anti-HLA-I antibody (1:20; clone W6/32, Invitrogen, 11-9983-42) or SNA-Cy5-lectin staining (1:100; VectorLabs, CL-1304-1;) and incubated for 30 min at 4°C, protected from light. Following centrifugation (400 × *g*, 5 min), cells were resuspended in FACS buffer containing Live/Dead Fixable Violet stain (1:1000; Invitrogen, L34694) and incubated for 20 min at 4°C. A live/dead staining control was prepared by heat-killing half of the unstained control sample (95°C, 5 min). Cells were washed twice in FACS buffer, resuspended in 400 µL FACS buffer, filtered through 35 µM strainer tubes (spl lifesciences, 40405). Acquisition was executed on a BD Fortessa. Cells were first gated on FSC-A vs. SSC-A to exclude debris, followed by singlet discrimination (FSC-H vs. FSC-A) and exclusion of dead cells using the Live/Dead Fixable Violet-negative population. Within the live, single-cell gate, HLA-I-FITC expression was assessed by histogram analysis to determine knockout efficiency, with all data analyzed using FlowJo v11 (BD Biosciences).

### Sanger Sequencing

Total nucleic acids were isolated from nasal basal cells by washing the cells with DPBS^-/-^, dissociating with 200 µL TrypLE for 5–10 min at 37°C and quenching with 300 µL AdDF+++, followed by centrifugation at 400 × *g* for 5 min. The cell pellet was washed once with AdDF+++ and resuspended in 90 µL MagNAPure lysis buffer (Roche, 06374913001). Total nucleic acids were subsequently extracted using 50 µL Sera-Mag SpeedBead carboxylate-modified magnetic particles per sample (Cytiva, 65152105050250), incubating at room temperature for 15 min, washing the beads thrice with 70% ethanol on a 96-well neodymium magnet. Nucleic acids were subsequently eluted in 50 µL DEPC-Treated Water (1^st^ Base, BUF-1170-1L) and concentration was determined using NanoDrop. The genomic region encompassing the CRISPR/Cas9 target site was amplified by PCR using pre-designed locus-specific forward and reverse primers and Takara HS Perfect Mix (Takara Bio, R300) according to manufacturer’s protocol. PCR products were assessed by electrophoresis on a 1% agarose gel, purified and sent alongside appropriate sequencing primers for Sanger Sequencing (BioBasic Asia). Sequencing chromatograms were subsequently compared between the gene specific and the non-targeting sgRNA control reference sequence to assess CRISPR/Cas9-induced KO efficiency using Inference of CRISPR Edits (ICE) analysis using the Synthego ICE web-based tool (Synthego).

### Western blotting

The apical surface of the nasal ALI cultures were briefly washed with DPBS^+/+^ (10 min, 37°C), before removing wash, removing the basal medium, and lysing in 60 µL Pierce™ IP lysis buffer (Thermo Fisher, 87788) supplemented with 1× Pierce™ protease inhibitor (A32955, Thermo Fisher) for 10 min on ice. Afterwards the lysate was collected by scraping and transferred to 1.5mL-eppendorf tube to rotate for 30 min at 4°C. Lysates were cleared by centrifugation (15,000 × *g*, 15 min, 4°C), and supernatants were mixed 1:1 with 2× Laemmli Sample Buffer (Axil Scientific, BUF-5240-25ml) and boiled for 20 min at 95°C prior to separation by SDS-PAGE. Proteins were resolved on 4–12% Tris-Glycine Plus WedgeWell gels (Invitrogen, XP04120BOX) using 1× TGS running buffer (Tris-base, Sigma-Aldrich, 93362-500G; Glycine, Sigma-Aldrich, 0000254755; 10% SDS, Invitrogen, 15553-035) at 50 V for 30 min, followed by 200 V for 40 min. Proteins were transferred to PVDF membranes (Roche, 03010040001) by wet transfer (25-30 V, 60 min, 4°C) in 1× transfer buffer containing 20% methanol. Membranes were blocked for 1 hour at room temperature in 5% skimmed milk or 0.5% BSA (Sigma-Aldrich, A9418-100G) in 0.1%Tween20-PBS (Tween 20, VWR, 0777-1L; DPBS^-/-^), then incubated with primary antibody; mouse anti-human TMPRSS2 (1:500; Abcam, ab309546) in 5% BSA, goat anti-human ACE2 antibody (1:200; R&D Systems, AF933) diluted in 5% milk or mouse anti-human GAPDH (1:1000; Sigma-Aldrich, MAB734) for 1-2 hours at room temperature or overnight at 4°C, followed by three 5-min washes at room temperature with 0.1%-Tween20-PBS on a rotor. Membranes were subsequently incubated with secondary donkey anti-goat/mouse/rabbit IgG Dylight™ 680 antibody (1:2000; Thermo Fisher, SA5-10170, SA5-10090, SA5-10044) or donkey anti mouse IgG Alexa Fluor™ Plus 555 (1:1000; Thermo Fisher, A32773) in blocking buffer for 60 min at room temperature, washed thrice with 5 min washes at room temperature in 0.1%-Tween20-PBS and imaged using a Li-Cor Odyssey scanner at 700/800 nm or or Invitrogen iBright for Alexa Fluor™ Plus 555. For TMPRSS2 detection by western blot, only the cleaved, active form of TMPRSS2 (∼25 kDa) was detectable, the full-length TMPRSS2 (∼55kDa) could not be detected in these cultures.

### Virus infection on differentiated nasal ALI organoids

Prior to infection, 6-week differentiated nasal ALI cultures were refreshed with 500 µL of fresh Pneumacult-ALI Medium. The nasal ALI cultures were washed once apically with 200 µL AdDF+++, incubated for 10 min at 37°C, before removal of the wash to eliminate residual surface mucus. After wash removal, the virus inoculum was prepared in AdDF+++ and added apically to each well at 1,000 FFU/well in 200 µL and incubated at 37°C for 2 hours. This corresponds to an approximate MOI of 0.01, based on an estimated 1 × 10⁵ luminally accessible cells and an MOI of 0.003, based on 300,000 total cells per 6.5mm Transwell ALI culture. After two hours of virus adsorption at 37°C, inoculum was harvested and cultures were washed thrice with excess AdDF+++ to remove unbound virus. Apical harvest was performed at designated hours post-infection time points (2, 24, 48, 72 h.p.i.), by adding 200 µL of AdDF+++ to the apical surface and incubating at 37°C for 10 min before collection by aspiration. Samples were stored at -80°C until processing.

### Viral titre quantification by RT-qPCR

As cell lines are not susceptible to clinical hCoV-NL63, we opted for RT-qPCR-based quantification instead of standard titration. In addition, we aim to use the CRISPR/Cas9 workflow to compare host dependency factors across SARS-CoV-2 variants, whose infectious titers can be underestimated in a variant- and cell type-dependent manner, whereas RT-qPCR-based quantification provides a consistent readout for comparisons across variants [33, 49]. For determining viral RNA titres, viral RNA was extracted from apical harvests by defrosting the samples at room temperature, mixing by pipetting and lysing 60 µL of apical wash in 90 µL of MagNA Pure lysis buffer for 5 min at room temperature. Viral RNA was purified using 50 µL Sera-Mag SpeedBead carboxylate-modified magnetic particles per sample and incubating at room temperature for 15 min, washing the beads thrice with 70% ethanol on a 96-well neodymium magnet. RNA was subsequently transferred to a fresh plate and eluted in 30 µL DEPC-Treated Water. Purified viral RNA was then quantified by one-step RT-qPCR targeting the SARS-CoV-2 envelope (E) gene using the AgPath-ID™ One-Step RT-PCR Kit (Thermo Fisher, 4387391) according to the manufacturer’s protocol, with the following primer-probe set for SARS-CoV-2 E [50]: Forward Primer ‘5 ACAGGTACGTTAATAGTTAATAGCGT 3’, Reverse Primer ‘5 ATATTGCAGCAGTACGCACACA 3’ and probe ’5-FAM-ACACTAGCCATCCTTACTGCGCTTCG-3’BHQ and for hCoV-NL63 N [51]: Forward Primer ‘5 GAAGCGTGTTCCTACCAGAGA 3’, Reverse Primer ‘5 GAATCCCCCATATTGTGATTAAA 3’ and probe ’5-FAM-AAATGTTATTCAGTGCTTTGGTCCTCGTGA-3’BHQ. Ct values were converted to copy numbers / µL per PCR reaction by interpolation from standard curves.

### Live-imaging confocal microscopy

Live imaging of infected nasal ALI cultures was performed using an Andor BC43 benchtop spinning disk confocal microscope equipped with an on-stage incubator to maintain culture conditions of 37°C and 5% CO_2_. Full-well images were acquired using a 10× objective by stitching 7×7 grids in brightfield, green-channel (488 nm excitation; 500–550 nm emission; eGFP) and yellow-channel fluorescence (561 nm excitation; 580–610 nm emission; mCherry). Images were analyzed using Imaris Spot analysis (version 11, Oxford Instruments) for spot/focus quantification, and ImageJ for particle counting and area-based quantification.

### Fixed-cell immunofluorescence confocal microscopy

ALI cultures were fixed in 10% Neutral-buffered formalin for 60 min, washed in PBS and stored in 4°C until staining or for crysections; cryoprotected in sequential 10%, 20%, and 30% sucrose (1st Base; BIO-1090-1kg) for 1 h at 4°C, followed by overnight equilibration in 30% sucrose/O.C.T. (Sakura; 4583). These samples were then embedded in O.C.T. and sectioned at 8 μm thickness using a cryotome. Sections were mounted on slides and stored at −80°C until staining.

Fixed nasal ALI cultures or their cryosections were permeabilized in 0.1% Triton X-100 in DPBS^⁻/⁻^ (0.1%-Triton-PBS) for 10 min and blocked in blocking buffer for 1 hour. Cells were incubated overnight at 4°C or 2 hours at room temperature with primary antibodies in blocking buffer, washed thrice with 0.1% Triton X-100 in DPBS^-/-^ and incubated with corresponding Alexa Fluor™ 488- or 555--conjugated secondary antibodies (1:1000; Invitrogen, A32794, A-21127, A32790) in blocking buffer supplemented with nuclei staining using Hoechst 33342 (1 µg/mL; Invitrogen, H3570) for 2 hours at room temperature protected from light. The samples were mounted in ProLong Gold antifade reagent (Invitrogen, P36934) and imaged on an Andor BC43 benchtop spinning disk confocal microscope. Confocal images were analyzed using Imaris 10.2.0 (Oxford Instruments).

Staining was performed for ciliated cells with Alexa Fluor™ 647-conjugated anti-acetylated-α-tubulin (1:500; clone 6-11B-1; Santa Cruz Biotechnology, sc-23950 AF647), goblet cells with mouse anti-MUC5AC (1:1000; clone 45M1; Invitrogen; MA5-12178), basal cells with rabbit anti-KRT5 (1:400; clone SC62-04, HuaBio; ET1610-43), SARS-CoV-2 infection with SARS-CoV-2 Rabbit anti-Nucleocapsid protein (1:1000; polyclonal; SinoBiological, 40143-T62), hCoV-NL63 infection with rabbit anti-hCoV-NL63 Nucleocapsid protein (1:1000, polyclonal, SinoBiological; 40641-T62).

### Entry route inhibitor assay

To assess the sensitivity of viral strains to entry pathway inhibitors, infections were performed as described above, with the following modifications. The apical compartment was pre-treated for 1 hour with 100 µM Camostat mesylate (MedChemExpress, HY-13512), 10 µM E64D/Aloxistatin (MedChemExpress; HY-100229), or an equivalent volume of DMSO (vehicle control) in AdDF+++. Washes and virus harvests were executed with AdDF+++ containing the respective inhibitor. RNA extraction and RT-qPCR were executed as described above.

### Statistical analysis

All statistical analyses were performed using GraphPad Prism 11.0.1 (GraphPad Software) with statistical details for each experiment provided in the corresponding figure legends.

### Use of artificial intelligence

AI tools (Claude Opus 5 and Claude Sonnet 5, Anthropic; ChatGPT GPT-5.6, OpenAI; Grammarly version 1.187.0.0, Grammarly Inc.) were used to check grammar and improve the structure and flow of existing text throughout the manuscript. All AI-assisted output was reviewed and edited by the authors, who take full responsibility for the content of the final manuscript.

## Acknowledgements

We thank Ze Ming Lim and the Duke-NUS Flow Cytometry Core Facility for assistance with sorting.

## Supplementary figures

**Supplementary Table 1.**
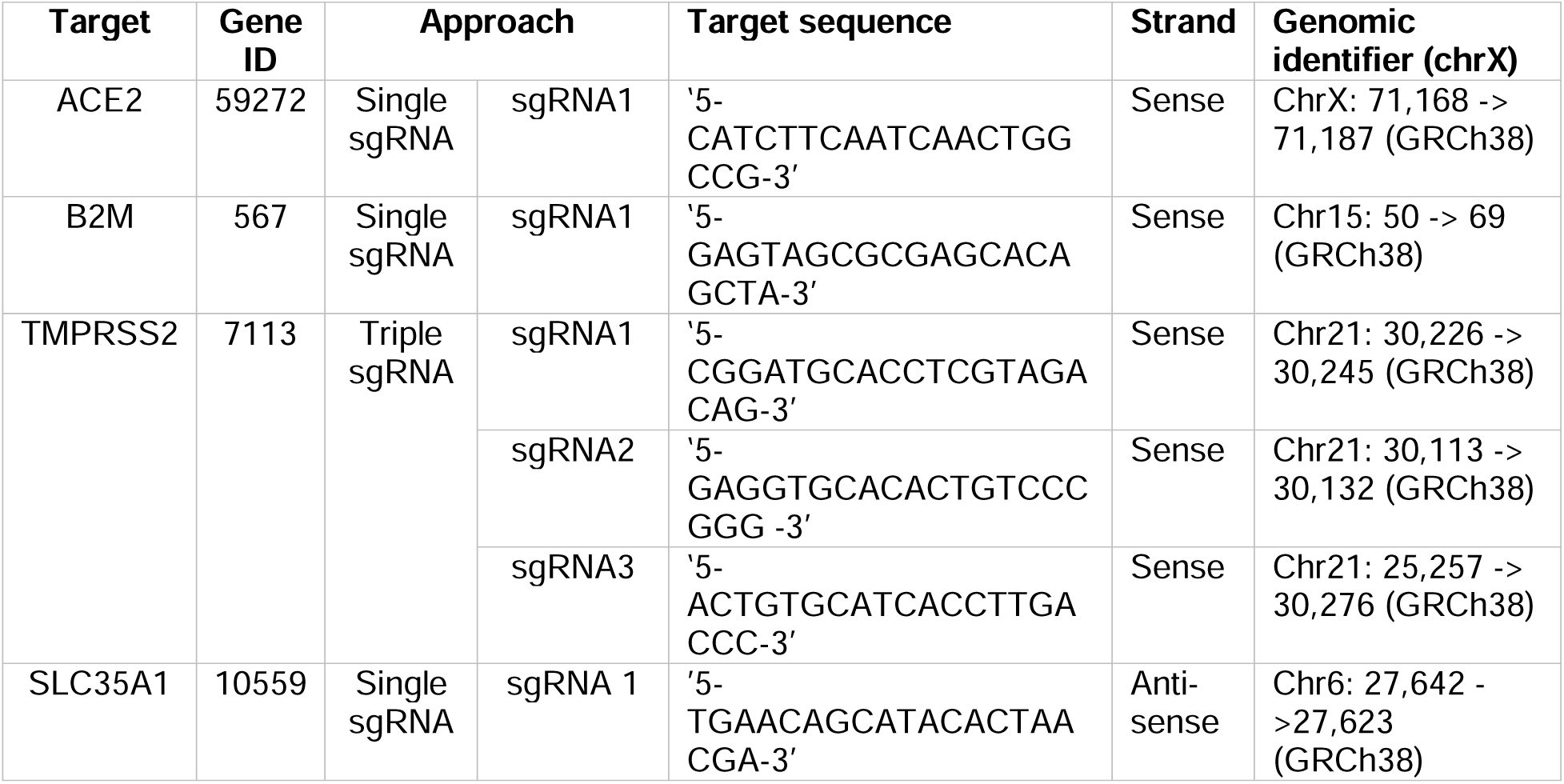
sgRNA sequences and information.

**Supplementary Table 2.**
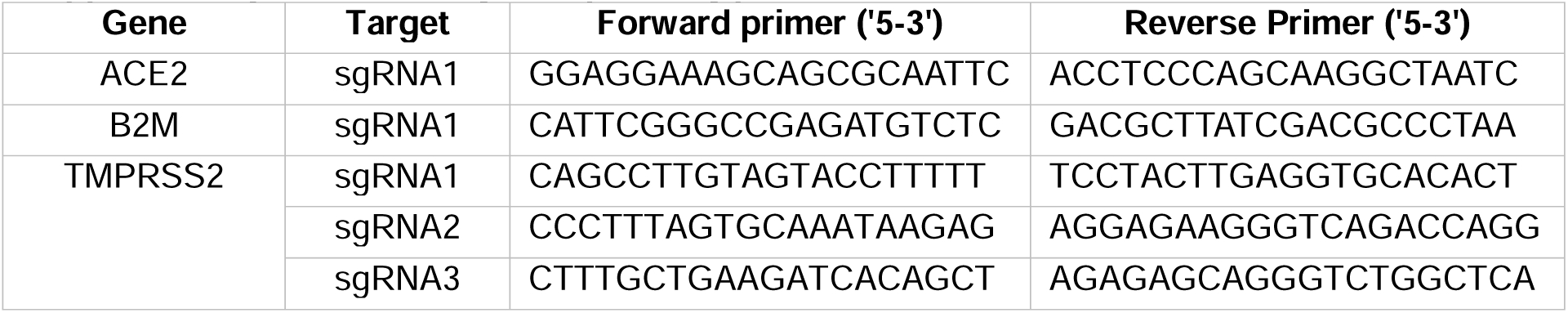
Sanger sequencing primers.

**Supplementary Fig. 1.**
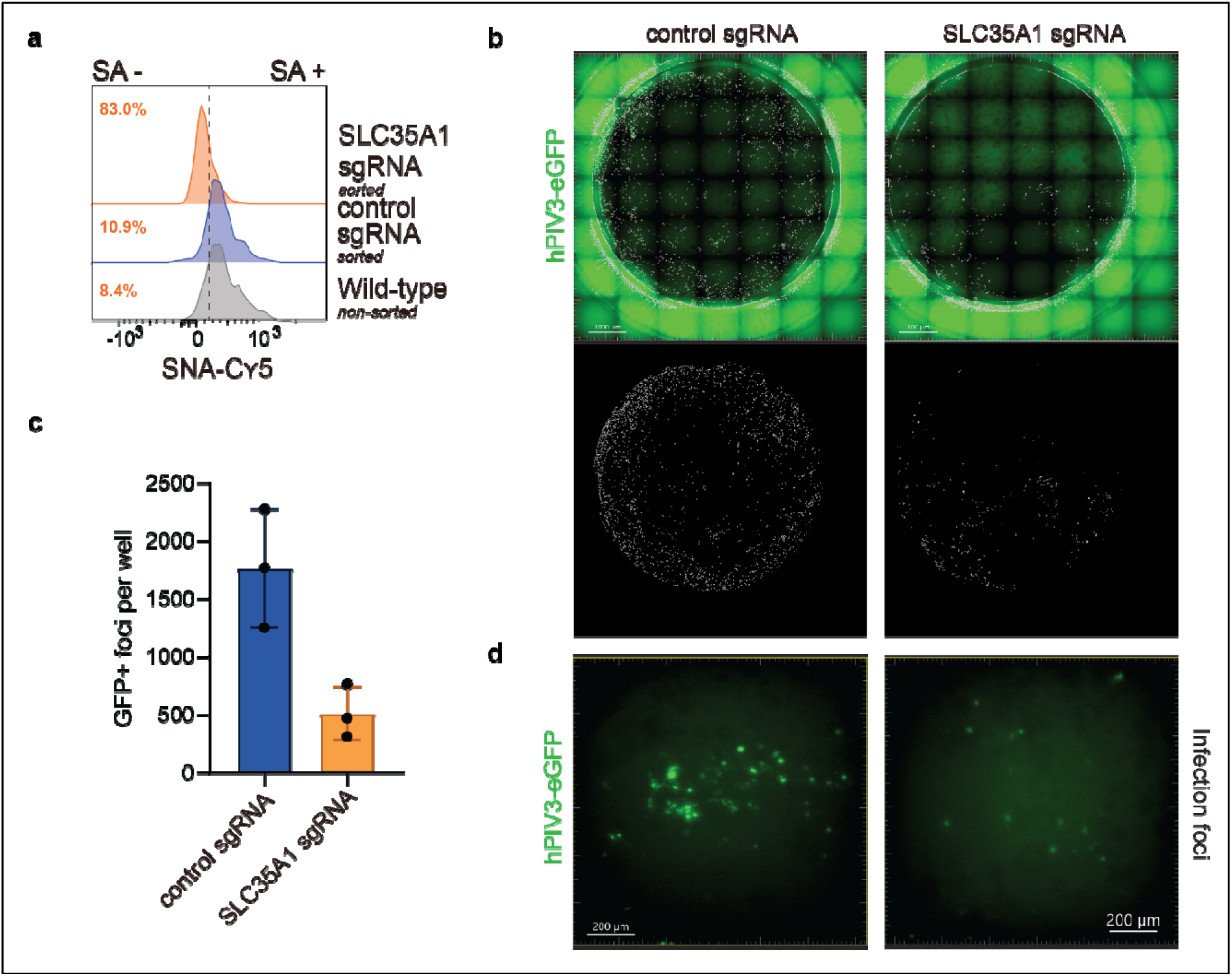
SLC35A1 knockout validates CRISPR/Cas9 workflow, with loss of terminal sialylation reducing susceptibility to hPIV3-eGFP. SLC35A1, encoding a sialic acid transporter required for cell surface glycosylation, was targeted as a proof-of-concept host factor given its established role in sialic acid-dependent viral entry, including hPIV3 [29]. The results presented here are representative of two independent SLC35A1 knockout batches. **a** Knockout of SLC35A1 abolishes terminal sialic acid (SA) expression, as measured by SNA-Cy5 staining, yielding an 83.0% SNA-negative population following GFP-based sorting. **b** Representative live-images of hPIV3-eGFP (100,000 FFU/well) infection in 3 weeks differentiated non-targeting sgRNA control vs. SLC35A1 knockout nasal ALI cultures at 72 hours post-infection. Images were acquired using a 10× objective and 7×7 grid stitching on an Andor BC43 benchtop spinning disk confocal microscope (488 nm excitation, green epifluorescence channel). hPIV3-positive foci were identified using Imaris 11 spot analysis (masks overlaid in white in **b** and quantified within a circular region of interest as defined in ImageJ, **c** confirming reduced hPIV3-eGFP infection in SLC35A1 knockout ALI cultures (triplicate ALI cultures, 1 donor). Each point represents one well; bars show mean ± SD. **d** Representative single 10× field (one grid position) from **b** that shows infection in both conditions, highlighting reduced eGFP signal intensity in SLC35A1 knockout compared to non-targeting sgRNA control ALI cultures, in addition to the reduced number of hPIV3-foci shown in **b** and **c**.

**Supplementary Fig. 2.**
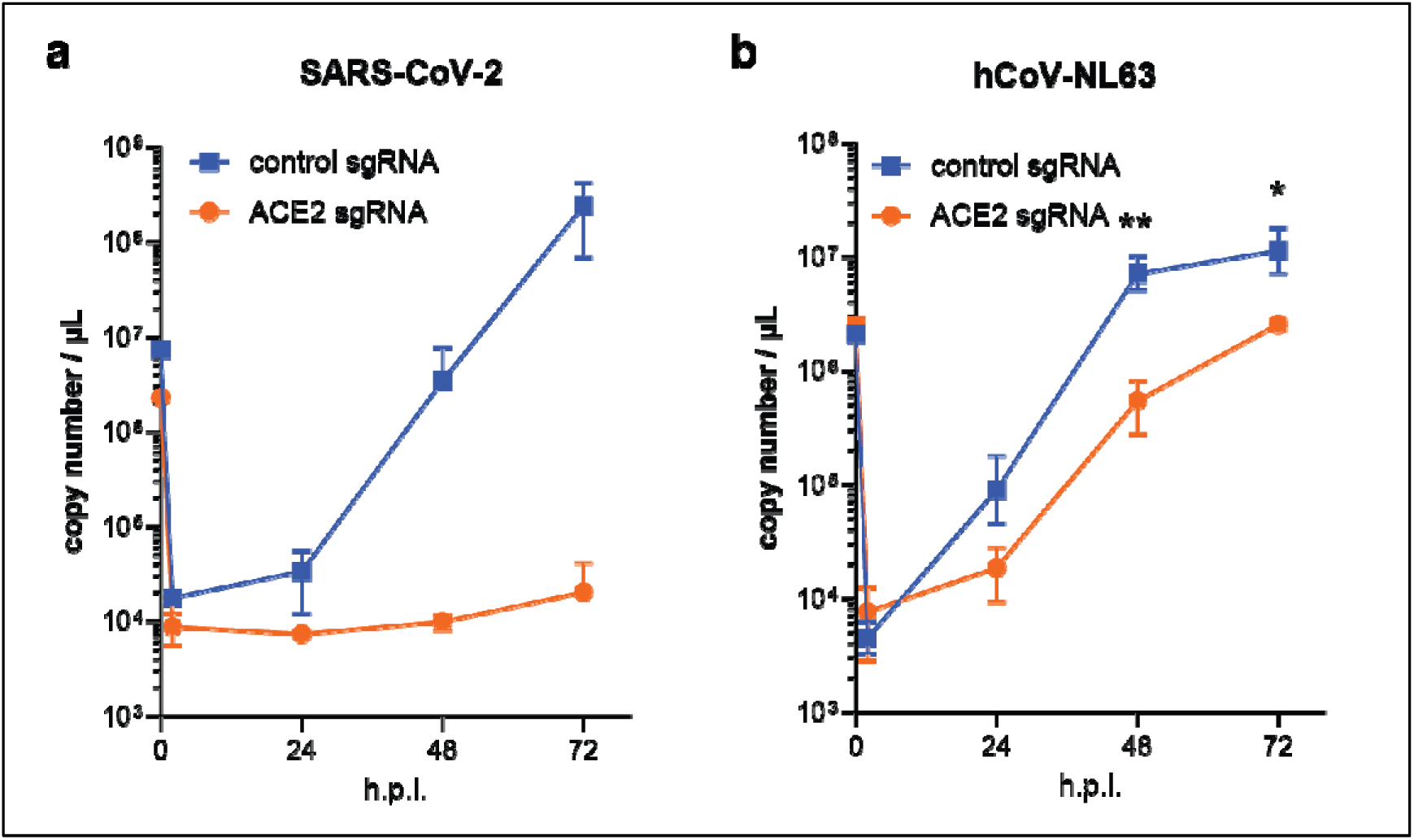
Independent ACE2 knockout batches confirm reduction in SARS-CoV-2 and hCoV-NL63 infection. **a** Nasal ALI cultures generated from an independent knockout batch using the same donor confirmed the loss of SARS-CoV-2 infection in the ACE2 sgRNA ALI cultures, as assessed by Envelope (E)-targeted RT-qPCR (duplicate ALI cultures per condition). **b** Triplicate ACE2 knockout (ACE2 sgRNA) ALI cultures generated in an independent donor background were infected with hCoV-NL63 at 1,000 FFU/well. Viral replication was quantified by nucleocapsid (N)-targeted RT-qPCR. Each data point represents the geometric mean of the **a** duplicate or **b** triplicate ALI cultures, with error bars indicating the geometric SD. Statistical significance was assessed if the experiment had at least 3 replicates using two-way ANOVA followed by Dunnet’s multiple-comparisons test. **b** TMPRSS2 and ACE2 sgRNA conditions were generated in the same knockout batch and therefore share the same non-targeting control data as the hCoV-NL63 experiment in **Fig. 3h**, comparing the knockout with the control at each time point with * p < 0.05 and ** p<0.01.

**Supplementary Fig. 3.**
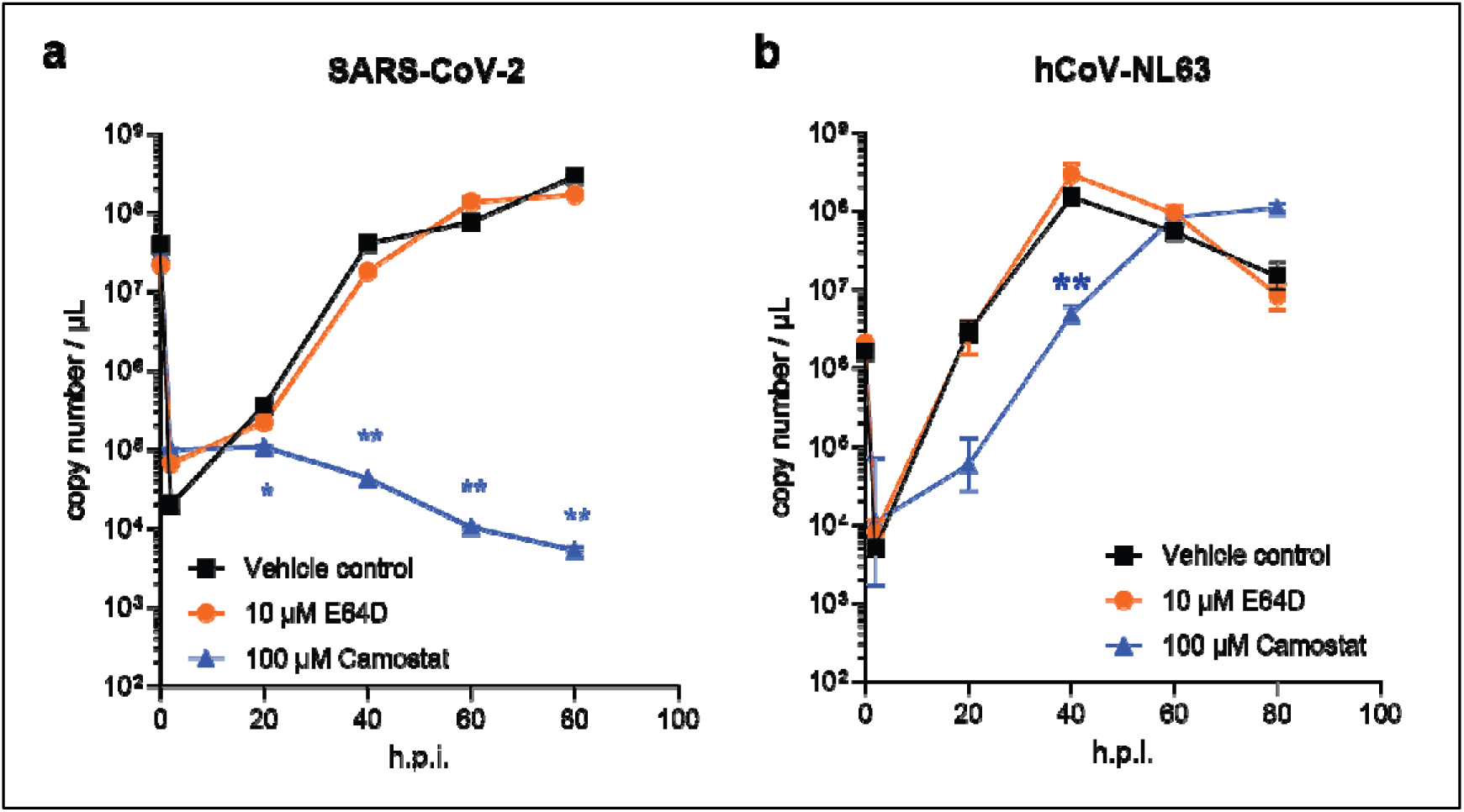
SARS-CoV-2 and hCoV-NL63 rely on serine protease-mediated, not cathepsin-mediated, entry in nasal ALI cultures. Results from independent donors confirm the inhibitor-based entry results. Nasal ALI cultures were infected with **a** SARS-CoV-2-mCherry (n = 1 donor, duplicate ALI cultures) or **b** hCoV-NL63 (n = 1 donor, triplicate ALI cultures) following apical pre-treatment with 10 µM E64D or 100 µM camostat. Viral infection was assessed by envelope (E) gene RT-qPCR for SARS-CoV-2 and nucleocapsid (N) gene RT-qPCR for hCoV-NL63 on apical washes collected at 2, 20, 40, 60, and 80 hours post-infection (h.p.i.), alongside the inoculum. Each data point represents the geometric mean of the duplicate or triplicate ALI cultures, with error bars indicating the geometric SD. Statistical significance was assessed using two-way ANOVA followed by Dunnett’s multiple-comparisons test, comparing each treatment or knockout with the control at each time point with * p < 0.05 and ** p<0.01.

**Supplementary Fig. 4.**
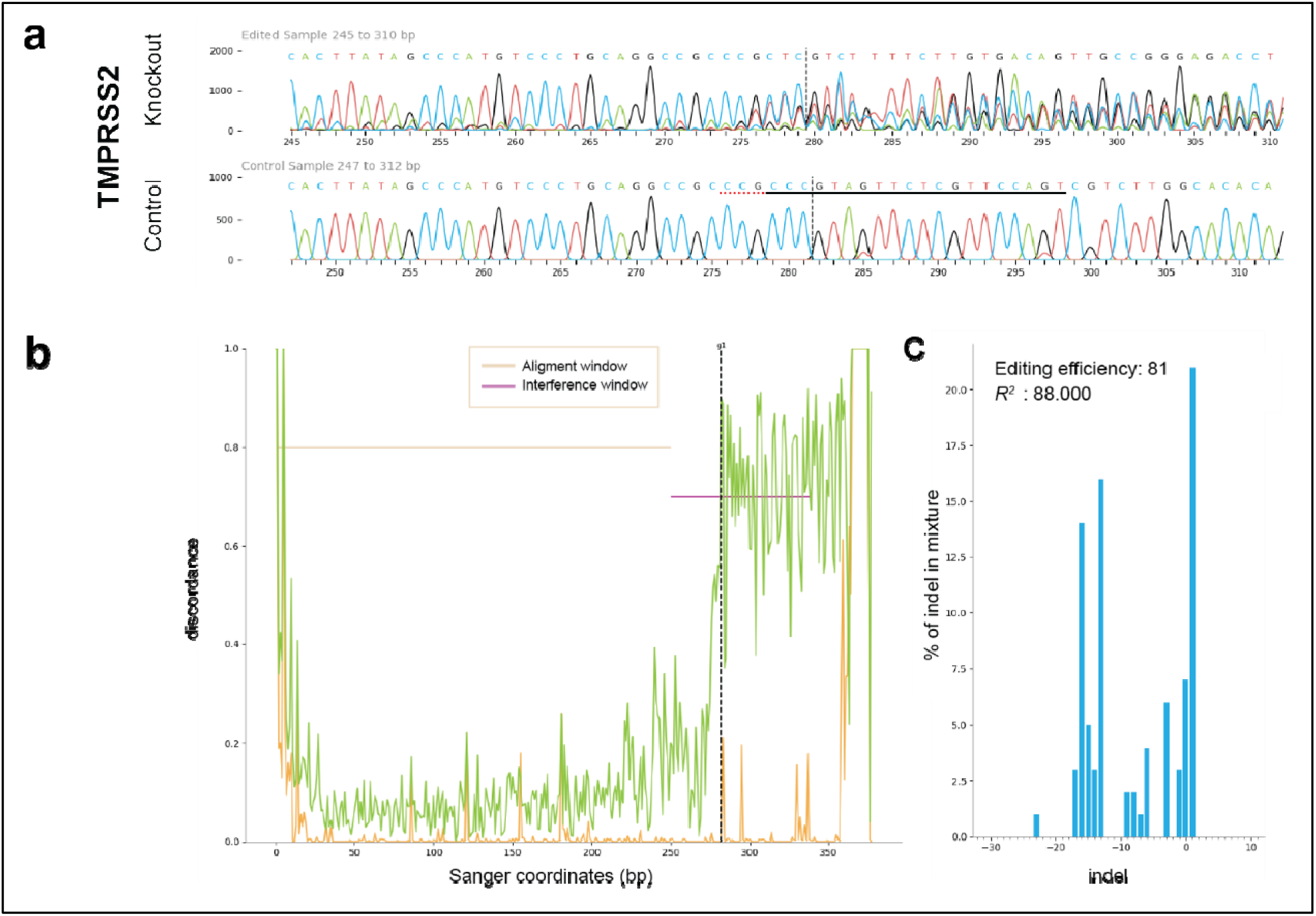
Representative Sanger sequencing traces from TMPRSS2-edited nasal basal stem cells following transfection and sorting. CRISPR/Cas9-induced editing efficiency was assessed using the Synthego ICE (Inference of CRISPR Edits) tool. **a** Sanger traces of the TMPRSS2 knockout (‘edited sample’) compared to a non-targeting sgRNA (‘control sample’), with the CRISPR/Cas9 cut site indicated by a dashed vertical line. **b** The discordance plot (left) shows a marked increase in sequence discordance downstream of the cut site. **c** The corresponding indel distribution shows the edited population is dominated by deletions (predominantly 10-20 bp), yielding an overall editing efficiency (ICE score) of 81%, a knockout score (frameshift-inducing indels) of 64%, an indel diversity (ICE-d) of 98%, and a model fit (R²) of 0.88, indicating high-confidence trace deconvolution.

**Supplementary Fig. 5.**
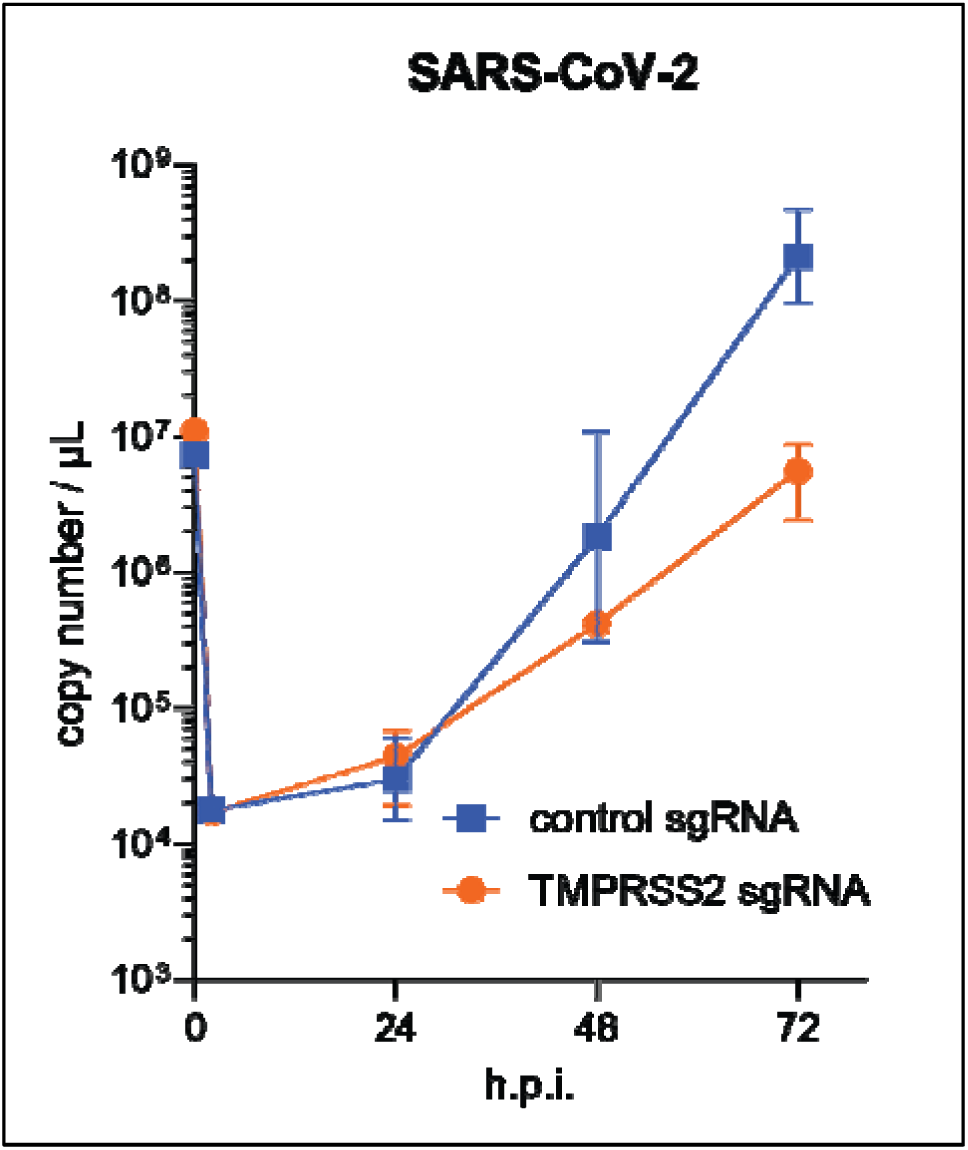
TMPRSS2 knockout confirmed to significantly reduce SARS-CoV-2 infection. Infection of nasal ALI cultures with SARS-CoV-2-mCherry at 1,000 FFU/well demonstrates significantly reduced viral RNA in an independent knockout nasal ALI culture batch for TMPRSS2 sgRNA in comparison to non-targeting control sgRNA. Viral replication was quantified by SARS-CoV-2 envelope (E)-targeted RT-qPCR of apical washes collected at 2, 24, 48, and 72 hours post-infection (h.p.i.); the input inoculum was analyzed in parallel. Each data point represents the geometric mean of two ALI cultures, with error bars indicating the geometric SD.

**Supplementary Fig. 6.**
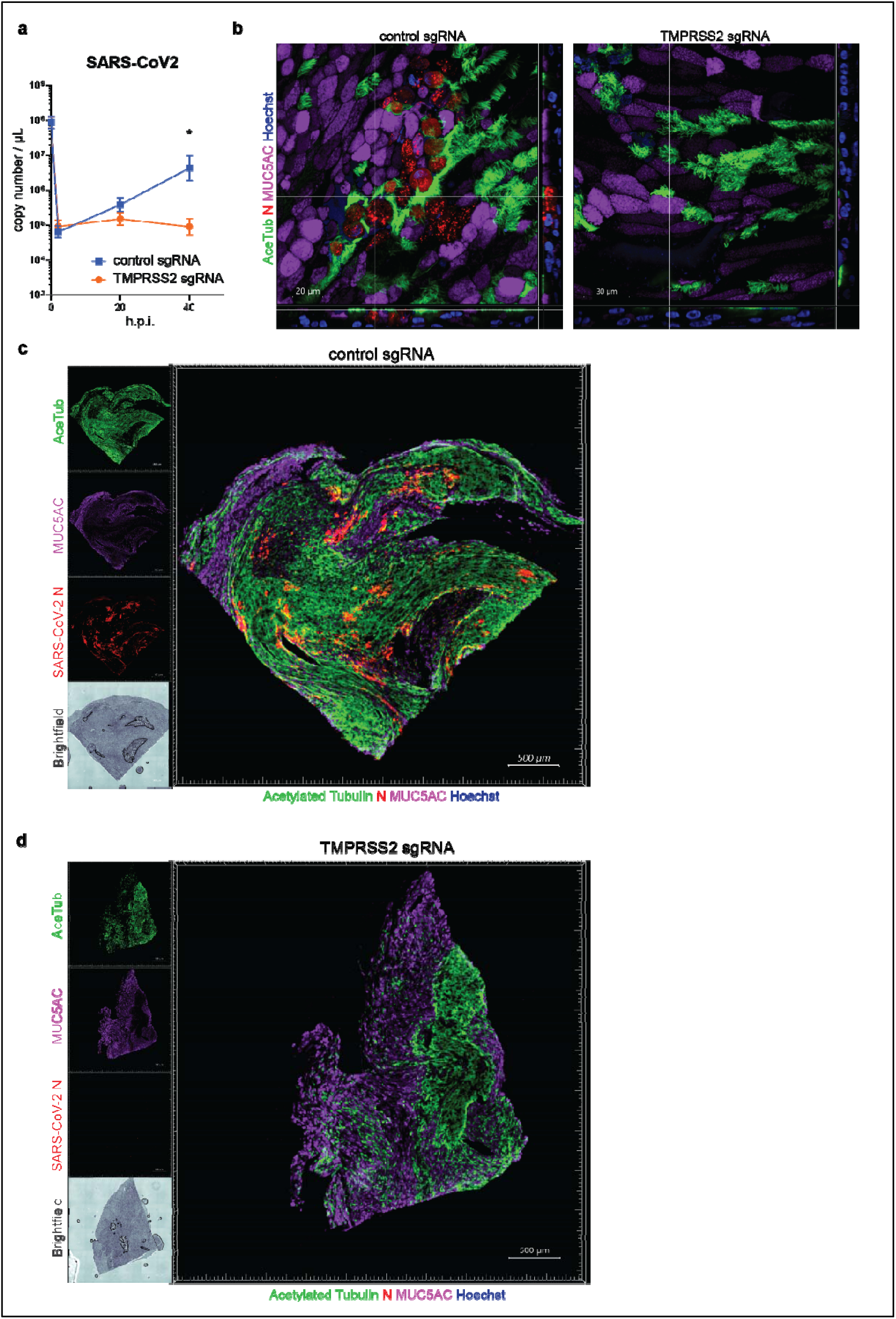
CRISPR-mediated TMPRSS2 knockout in a second donor abrogates SARS-CoV-2 infection. **a** TMPRSS2 knockout in an alternative donor shows abrogated SARS-CoV-2-mCherry infection, as measured by RT-qPCR (E gene RT-qPCR; n = 1 donor, triplicate ALI cultures). Each data point represents the geometric mean of the triplicate ALI cultures, with error bars indicating the geometric SD. **b** Differentiated ALI cultures retain ciliated (Acetylated tubulin+ (AceTub)) and goblet (MUC5AC+) cells in both triple-guided TMPRSS2 knockout (sgRNA TMPRSS2) and non-targeting control sgRNA (control sgRNA) conditions (confocal, 40×), but **c-d** Immunofluorescence staining of quarter cut-outs of fixed samples at 40 hours post-infection (h.p.i.) shows loss of infection in TMPRSS2 sgRNA, evidenced by the loss of SARS-CoV-2 nucleocapsid protein (N) staining, compared to widespread infection in non-targeting sgRNA ALI cultures. Statistical significance was assessed using two-way ANOVA followed by Holm-Šidák’s multiple-comparisons test, comparing each treatment with the control at each time point with * p < 0.05.

